# Modeling of chemo-radiotherapy targeting growing vascular tumors: a continuum-level approach

**DOI:** 10.1101/2024.03.21.586183

**Authors:** Ioannis Lampropoulos, Marina Koutsi, Michail Kavousanakis

## Abstract

The aim of this study is to demonstrate the enhanced efficiency of combined therapeutic strategies for the treatment of growing tumors, based on computational experiments of a continuous-level modeling framework. In particular, the tumor growth is simulated within a contaminated tissue and treated as a multiphase fluid of high viscosity, with each cellular species considered as a distinct fluid phase. Our model integrates the impact of chemical species on tumor dynamics, and we model –through reaction-diffusion equations– the spatio-temporal evolution of oxygen, vascular endothelial growth factor (VEGF) and chemotherapeutic agents. Simulations of a growing tumor exposed to external radiation showcase the rapid impact of radiotherapy on tumor suppression, however this effect diminishes over time. To enhance the therapeutic efficiency of radiotherapy, we investigate the combination of external radiation with the anti-VEGF drug bevacizumab and the cytotoxic drug docetaxel. Our simulations demonstrate that this synergistic approach integrates the immediate effectiveness of radiation therapy with the enduring tumor-suppressive capabilities of chemotherapy.

## Introduction

Cancer, a pervasive and intricate disease, can manifest throughout the human body and metastasize [1]. Given its severity and complexity, recent clinical practices often employ a combination of therapies to treat patients. In 1965, Frei et al. conducted a pioneering clinical trial that combined chemotherapeutic agents in treating children with acute lymphocytic leukemia [2, 3]. Since then, ongoing research and clinical trials continuously explore and refine combination therapies to optimize cancer treatment strategies. This blended approach seeks to maximise therapeutic efficacy while reducing the risk of drug resistance emerging within the tumor [2, 4].

Radiotherapy, a cornerstone of many treatment regimens, is estimated to be administered to over half of all cancer patients during their course of treatment [5]. It employs precisely focused, high-energy beams to destroy tumors and prevent their recurrence. The most common mechanism of radiation-induced cell death for solid tumors is mitotic death [6, 7], where irradiated cells undergo one, two, or a few divisions before succumbing hours or days later [6, 8, 9]. Baskar et al. [10] and Lee et al. [11] underscore the pivotal role of radiotherapy, particularly as definitive treatment for certain cancers (e.g., prostate cancer). Various administration methods, including external beam radiation therapy and brachytherapy (internal radiation therapy, where radioactive material is delivered in the body near the infected site) [12, 13] are employed in clinical practice. The dose and duration of therapeutic sessions vary based on factors such as the tumor size, radiation sensitivity, and surrounding tissue resilience [12, 14].

In conjunction with radiotherapy, conventional clinical practice incorporates the administration of chemical drugs designed to either eliminate cancer cells or target specific molecular components. Cytotoxic chemotherapy stands out as one of the oldest and most prevalent tools in the fight against cancer. This therapeutic approach not only achieves the eradication of tumors but also augments the prospects of successful surgical removals with complementary methods such as radiotherapy [15]. Among the cytotoxic drugs, taxanes form a family extensively employed in various cancers, including ovarian and breast cancer. In particular, taxanes, exert their effects by targeting microtubules within cancer cells. This mechanism involves stabilizing these structures and impeding their depolymerisation - a pivotal step in mitotic cell division. Consequently, this blockade disrupts the mitotic process, ultimately inducing apoptosis [16, 17]. It is noteworthy that docetaxel (brand name Taxotere), as a semi-synthetic member of the taxanes class of medications, is included in W.H.O.’s List of Essential Medicines [18].

Various chemical agents offer alternative strategies to impede cancer growth.

Anti-angiogenesis drugs, instead of directly inhibiting mitosis and inducing immediate cell death, focus on disrupting the tumor’s vital supply chain. Specifically, these drugs thwart the formation of a peritumoral vascular network by binding to relevant growth hormones [19, 20]. Bevacizumab, commercially known as Avastin, specifically targets the vascular endothelial growth factor (VEGF), which is essential for the survival of proliferating endothelial cells, leading to their regression (in the absence of VEGF) [19, 21–23].

To improve the delivery of radiotherapeutic and chemotherapeutic agents, many clinical trials and investigations have been made regarding combining treatment modalities. Current studies examine the effects of new combination treatments, especially the treatment modality that combines radiation, and the agents of taxane, and bevacizumab [24]. Hainsworth et al. [25] performed a phase II trial of combined modality treatment with chemotherapy, radiation therapy, bevacizumab, and erlotinib in patients with locally advanced squamous carcinoma of the head and neck. Another pilot trial was performed by Wozniak et al. [26] regarding the incorporation of bevacizumab with concurrent radiotherapy in the treatment of locally advanced stage III non-small-cell lung cancer (NSCLC). Apart from clinical trials, radiotherapy is combined with taxane chemotherapy in clinical practice, with the LULACATRT protocol for non-small cell lung cancer [27] and the GIENACTRT protocol for esophageal and gastroesophageal carcinomas [28]. On the other hand, docetaxel and bevacizumab is also a common combination in several cancer treatments [29–31].

Radiation therapy and chemotherapy have been explored through both experiments and computational modeling. Several studies have focused on modeling the impact of radiotherapy on cancerous tumors, with the linear-quadratic (LQ) model proposed by Fowler [32] standing out as the most well-known and straightforward formula. This model predicts the surviving fraction of tumor cells within an irradiated tumor and has served as a foundational framework for various modeling approaches [33–37]. Enderling et al. [38] provided comprehensive overview of common computational approaches to simulate radiotherapy.

Several computational approaches have been developed for the study of the impact of radiotherapy, cytotoxic chemotherapy, anti-angiogenetic therapy and their combinations on growing tumors [39–45]. Kirkby et al. [46] developed a population balance model for an individual treated with radiation therapy for glioblastoma, later extending it to a population-based model using Monte Carlo. This extension was further advanced by Barazzuol et al. [33] three years later to investigate the potential benefits of temozolomide in the context of of chemo-radiotherapy combinations or as an adjuvant therapy. Their study suggests that temozolomide may enhance the tissue’s sensitivity to radiation therapy.

Moreover, several mathematical models employing partial differential equations (PDEs) have been developed to elucidate relevant phenomena and spatial population distributions within complex systems. Among these models we report the study of Enderling et al. [34] simulating single-fraction radiotherapy for breast cancer. Powathil et al. [47] devised a generalized LQ model to simulate various radiotherapy schemes for gliomas. Kohandel et al. [48] explored the combined effects of cytotoxic and anti-VEGF drugs in a two-dimensional model. Lastly, Mi et al. [36] proposed a system of PDEs to monitor the expansion of lung cancer under radiotherapy.

In the present study, we build upon existing literature [49–52], extending our PDE-based model to incorporate the impact of external radiation therapy. Our radiotherapy model posits that the death rate of tumor cells induced by radiation follows and exponentially decaying function over time with a specific half-life. Expanding beyond radiation monotherapy, we enhance our external radiotherapy model by introducing cytotoxic and anti-VEGF chemotherapeutic drugs as part of a combination therapy within a two-dimensional domain. To the best of our knowledge, this is the first time a unified computational model integrating the impact of radiation, cytotoxic chemotherapy and anti-angiogenic therapy on a growing vascular tumor is presented. Lastly, we delve into the phenomenon of radiosensitization during which, specific agents have the ability to amplify the efficacy of radiotherapy when combined [53]. More specifically, we are looking into taxanes induced radiosensitization [54], as well as oxygen as a radiosensitizer [55]. Our model is solved using the commercial software Comsol Multiphysics, and computes the spatio-temporal evolution of cellular and chemical species in an infected two-dimensional model tissue.

The paper is structured as follows: in Section Models, we describe the mathematical formulation of our model, focusing on the implementation/integration of radiotherapy in the proposed computational framework. Section Results presents simulations elucidating the impact of standalone radiation treatments and the combined approach of radio-chemotherapy on tumor growth and expansion. Section Conclusion encapsulates a summary of our results, a discussion on the merits and drawbacks inherent in this modeling approach, and recommendations for future research.

## Models

In this section, we consolidate the basic mathematical relationships that underpin our two-dimensional multiphase model. We conceptualize the contaminated tissue as a fluid composed of distinct, non-miscible, and interacting cellular-liquid phases. These phases include: I. Healthy cells, denoted with *h*. II. Cancer cells, denoted as *c*, forming the building blocks of the malignant tumor. III. Young (immature) blood vessels, represented with *yv*, that are products of pathological angiogenesis. IV. Mature blood vessels, denoted as *mv*, shaping the tissue’s capillary network. V. Interstitial fluid, represented with *int*, containing water and solutes.

In addition to cellular phases, the presented model considers four distinct chemical species integral to the described phenomena. These chemicals include: I. Oxygen, which represents nutrients and is denoted with, *c*. II. The vascular endothelial growth factor (VEGF), denoted with *g*, a principal factor in angiogenesis and peritumoral vascular network formation. III. The anti-angiogenesis drug bevacizumab, represented with *a*, which inhibits VEGF. IV. The cytotoxic drug docetaxel, denoted with *w*, which belongs to the taxane medications family, and exhibits cytotoxic properties.

The model’s core elements encompass mass balance equations for both fluid/cellular and chemical species, as well momentum balance equations employed to compute the macroscopic flow of each fluid phase. Momentum balance equations are complemented by auxiliary algebraic expressions, as detailed bellow.

### Cellular Phases

The volume fraction of each cellular phase, *i*, considered in the model is computed through the solution of their mass balance equations. In addition, each phase possesses its velocity field, denoted with 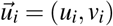, and pressure, *p*_*i*_. The velocity fields are computed by solving momentum balance equations for compressible fluids in a creeping flow, while pressures are determined through the continuity equation (mass balance for all cellular species) complemented by algebraic equations of state.

### Mass balance equations for cellular phases

It is reasonable to assume a uniform density for the tissue being modeled at the spatial scales we are considering. Following this assumption, the mass balance equation can be formulated for each cellular phase (healthy and cancer cells, mature vessels, and interstitial fluid) as follows:

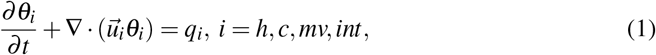

where *θ*_*i*_ denotes the volume fraction of cellular phase, *i*. The term *q*_*i*_ denotes the source term for the cellular species, and encompasses all processes related to mass transfer between them.

While the primary transport mechanism for healthy cells, cancer cells, mature vessels, and interstitial fluid is convection (the second term in Eq. (1)), for young vessels we introduce chemotaxis as an additional mechanism. In particular, the mass balance equation for young vessels reads:

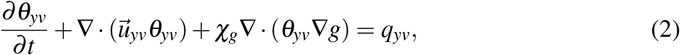

where χ_*g*_ denotes the chemoattraction potency parameter. The vascular network in healthy tissues is fully developed and primarily consists of endothelial cells [56]. The endothelial cells at the vascular stalk’s tip exhibit sensitivity to chemotaxis, driven by the presence of vascular endothelial growth factor (VEGF), leading to directed migration towards regions with higher VEGF, *g*, concentrations [57].

A fundamental premise of our model is based on the assumption that the tissue is a closed system, ensuring zero net mass transfer between cellular species (a common assumption of a wide selection of relevant models [52, 58–63]):

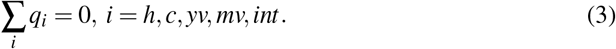

Lastly, the no-void assumption is incorporated:

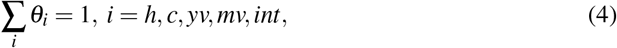

indicating that the tissue domain exclusively comprises the cellular/fluid phase components outlined in the model.

### Momentum balance equations for cellular phases

It is crucial to emphasize that the primary mode of cellular species transportation is convection. To delve deeper into this process, it becomes essential to recognize that the cellular movement is more accurately characterized by creeping flow, a low Reynolds number flow. This approach allows the formulation of momentum balance equations capturing the intricate dynamics of cellular species transportation:

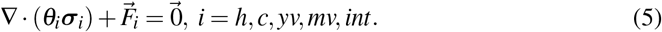

Here, the stress tensor, denoted by *σ*_*i*_, represents that of a high viscosity, compressible fluid.

In addition, it is important to highlight that computing of pressure fields, *p*_*i*_, for each cellular phase, involves employing carefully selected algebraic equations, commonly referred to as closures. One such equation, integrated into our model, originates from the assumption of a closed system. This assumption allows us to sum up the mass balance equations of all cellular species, Eq. (1)-(2), and incorporate Eq. (3) into the analysis. This approach ensures proper consideration of various components, providing a thorough understanding of pressure fields and their impact on the overall dynamics of the system:

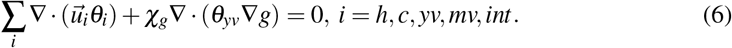

Completing the set of algebraic equations for pressure fields involves a specific condition where vascular phase pressures, *p*_*yv*_ and *p*_*mv*_, are equated to a reference pressure, *p*_*ref*_ . This reference pressure, which is exerted onto the tissue, is set at zero (*p*_*ref*_ = 0). The algebraic constraints for pressure fields of healthy and cancer cells follow the work of Hubbard and Byrne [49]:

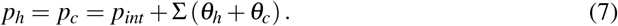

where Σ(*θ*) becomes non-zero, when the cellular density *θ* surpasses the numerical value, *θ*^*^, it maintains within a healthy tissue:

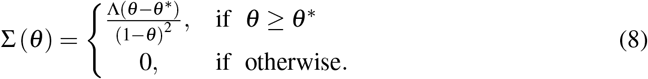

Here, Λ is a tension constant measuring the tendency of cells to restore their natural density.

### Chemical Species

In our modeling approach, we have opted to represent nutrient chemical species by focusing on oxygen. While the significance of the nutrient, *c*, and VEGF, *g*, remains paramount as inherent components of the studied ecosystem, the inclusion of anti-angiogenesis drug (bevacizumab), and cytotoxic drug (docetaxel) is pivotal for the advancement of our model extension. This extension enables us to investigate the intricate interplay of radiation with anti-VEGF chemotherapy, radiation with cytotoxic chemotherapy, as well as the combined effects of all therapies. The chemical species under consideration are assumed to be soluble in all fluid phases, and their relatively small size renders any contribution to the fluid volume negligibly small. Additionally, the dynamics of these chemical species are deemed to be significantly rapid compared to the cellular phases, enabling the adoption of a quasi-steady state approach.

#### Mass balance equations for chemical species

The diffusion is recognized as the primary mechanism governing the transportation of chemical species; considering their rapid dynamics, we solve the associated mass balance equations in steady-state conditions. Consequently, the general equation governing mass transfer for chemical species can be formulated as:

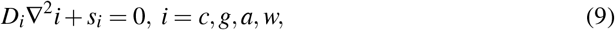

where, *D*_*i*_ denotes the diffusion coefficient of chemical species, *i*, and *s*_*i*_ represents the corresponding source term associated with said species. The mass balance equations for all chemical species incorporated in our model are complemented by zero-flux (Neumann) boundary conditions.

### Therapy-free model and parameters

For a complete description of the model and a comprehensive listing of parameters pertinent to the therapy-free model, we refer the reader to the Supplementary Material of the the manuscript, chapters II and III. We also refer the readers to previous works [50, 51] for a more in-depth analysis of both the therapy-free and the combination chemotherapy model. In the present study, we concentrate on novel enhancements introduced, specifically addressing adaptations necessary for incorporating radiation within our model, particularly concerning the cancerous cells phase. In light of this particular rationale, it is worth noting that the current research abstains from presenting any form of source terminology related to equilibrium equations for diverse cellular and chemical entities, as this is not the central focus of the present investigation.

### Radiotherapy, anti-VEGF and cytotoxic chemotherapy

The integration of radiation is a significant expansion to our established model, as elaborated by Lampropoulos et al. [50, 51]. We incorporate both standalone radiation therapy and combined radio-chemotherapy, involving two drugs: the anti-VEGF drug bevacizumab and the cytotoxic drug, docetaxel. Radiation administration involves the precise application of an external beam, targeting the affected region of the malignant growth in the patient’s organism [14].

External radiation therapy often divides the total dose into smaller fractions, typically administered over several weeks to minimize collateral damage to normal tissues. Each therapeutic session typically lasts approximately 30 to 45 minutes, with treatments administered five days a week and weekend breaks, spanning five to eight weeks [64]. Conventional fractionation treatment in radiotherapy employs fraction sizes of 1.8 to 2 *Gy* resulting in a total weekly dose of 9 to 10 *Gy* [65]. When used as adjuvant therapy, external beam radiation doses typically range from 45 to 60 *Gy* for various cancer types such as breast, head, and neck cancers [64].

On the other hand, both pharmaceutical agents are introduced into the human body through intravenous administration [66] and eventually reach the infected tissue through the vascular network. The transportation of drugs within the tissue occurs through diffusion.

#### Radiation

In the present study, we propose that radiation exclusively impacts the cellular phase of cancer cells. In particular, Eq. (10) models the cancer cell death rate due to radiotherapy [52], and formulates an exponential decay over time, with a half-life of 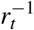 for successive radiation fractions administered at preset intervals:

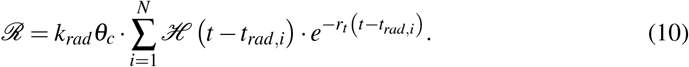

Here, the parameter *k*_*rad*_ quantifies the strength of radiotherapy dose, 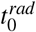 is the time of the first radiation, and *t*_*rad,i*_ signifies the initiation time of each radiotherapy session, *i*, ranging from *i* = 1 to *N*. ℋ denotes the Heaviside function activating each radiotherapy session for times, *t* > *t*_*rad,i*_. The parameter *r*_*t*_ indicates the half-life associated with the death of tumor cells due to radiotherapy. Finally, the time of the *i*^*th*^ radiation administration, *t*_*rad,i*_, is calculated using:

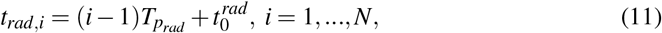

with 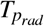 the period between two radiotherapy sessions.

Incorporating radiation in our model involves modifying the cancer cell source term, *q*_*c*_, to accommodate the radiation-induced cellular death kinetics described in Eq. (10):

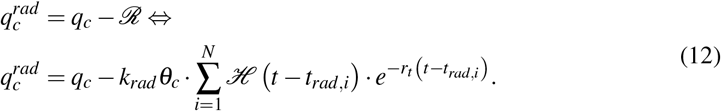

### Radiation therapy parameter determination

In this section, we provide details for the determination of parameter values used for the implementation of our radiotherapy model. We start our analysis by re-visiting the LQ model that serves as a pivotal mathematical formulation quantifying the survival fraction (SF) of cell colonies under specific radiation doses. LQ provides a practical and straightforward means of establishing the relationship between SF and radiation dose, taking the form of an exponential function encompassing both linear and quadratic components [67, 68]. The pioneering work of Fowler [32] involved calculating the SF for individual cells at a specific dose level, denoted as *d*, utilizing the LQ cell survival model:

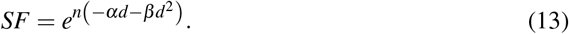

Here *n* represents the number of fractions, while *d* denotes the energy absorbed for each individual dose fraction (measured in *Gy*). Consequently, the product of *n×d* signifies the overall dose measured in Grays. Furthermore, *α* and *β*, represent the linear and quadratic coefficients, respectively. Notably, the values of *α* and *β* are intrinsically linked to the radio-sensitivity of cells. For numerous tumors, the ratio 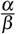 can be estimated from empirical observations, providing valuable insights into treatment [69].

In our simulations, numerical values for *α* and *β* are assigned based on the intrinsic radio-sensitivity of the cell type being modeled [70]. In particular, we utilize the widely adopted ratio [71]:

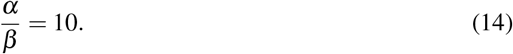

This choice is supported by the LQ model’s ability to accurately replicate in vitro cell survival and depict the impact of clinical fractionation, offering researchers and medical professionals a consistent and reliable means of interpreting the effects of fractionation on tissues and tumors [72, 73].

In a study by Higashi et al. [74], an irradiation dose of 30 Grays for a single fraction resulted in a significant and exponential decrease in viable cells, emphasizing the profound impact of radiotherapy on cellular viability. This reduction can be quantified by the SF of the cells, which serves as a metric for evaluating radiotherapy efficacy.

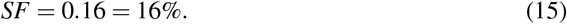

Then, we can derive values for the parameters *α* and *β* solving the system:

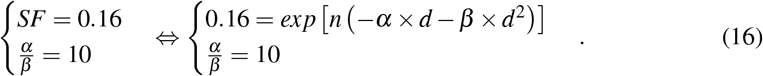

Since the clinical study provides information on the dosage of irradiation administered, the values of *n* = 1 and *d* = 30 Grays are set [74] to compute:

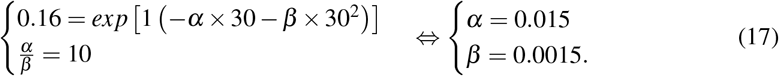

Assuming cancer cells are evenly spread throughout the body, we explore the potential impact of radiotherapy on the cell death rate. The term for cancer cell death caused by radiotherapy Eq. (10) can be formulated as:

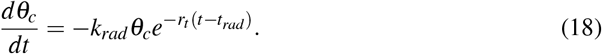

Integrating Eq. (18):

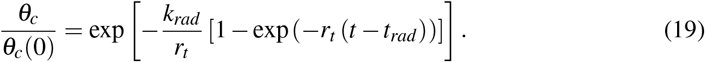

Based on the findings of Higashi [74], tumor cells experienced a decline of 84% over the course of a span of 12 days. This implies, that we can adjust the values of parameters, *k*_*rad*_ and *r*_*t*_, which gives 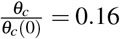 when *t* = 12. The fitting process produces the following set of parameter values: *r*_*t*_ = 0.5 and *k*_*rad*_ = 0.91, and in Fig. 1 we illustrate the evolution of SF according to Eq. 19. In our model, the parameter *r*_*t*_, denoting the half-life of cell death induced by radiotherapy, remains constant as it is contingent upon the intricate interplay between tumor cells. The *k*_*rad*_ value quantifies the radiotherapy intensity, and can be adjusted based on the radiation dosage administered in each treatment session, allowing for modifications in case of a different radiotherapy schedule.

**Fig 1.**
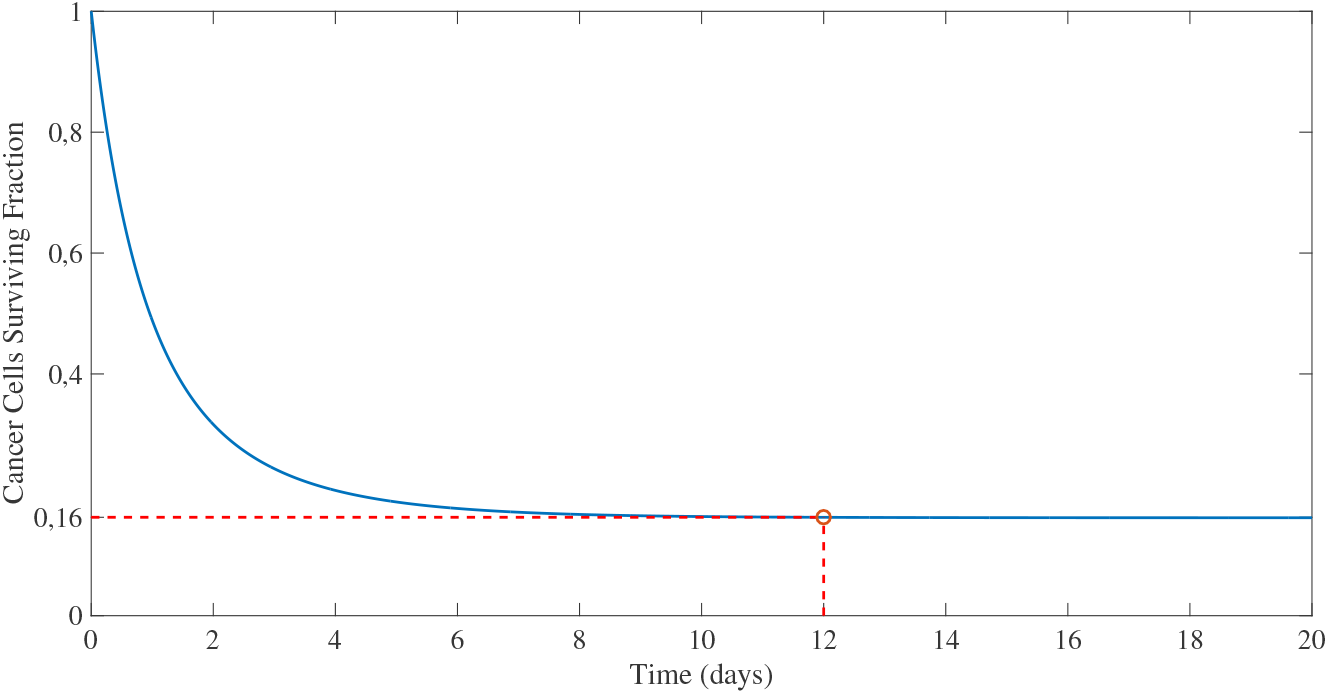
Evolution of viable tumor cells after irradiation according to Eq. (19).

More specifically, in the case of 30 fractions of 2 *Gy*, the LQ model predicts a *SF* ≈ 34%. Through trial and error, we compute that *k*_*rad*_ = 0.23 leads to a tumor that at the end of the therapeutic course, has a surviving fraction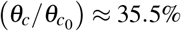. The deviation between LQ and our model is approximately 4.6%, and can be considered as acceptable, given that in our model, the surviving cells undergo several mitosis cycles during this time interval (the LQ model does not account for mitosis cycles). Hence, the value *k*_*rad*_ = 0.23 was selected for our simulations.

Table 1 summarises the parameters relevant to the implementation of radiotherapy.

**Table 1.**
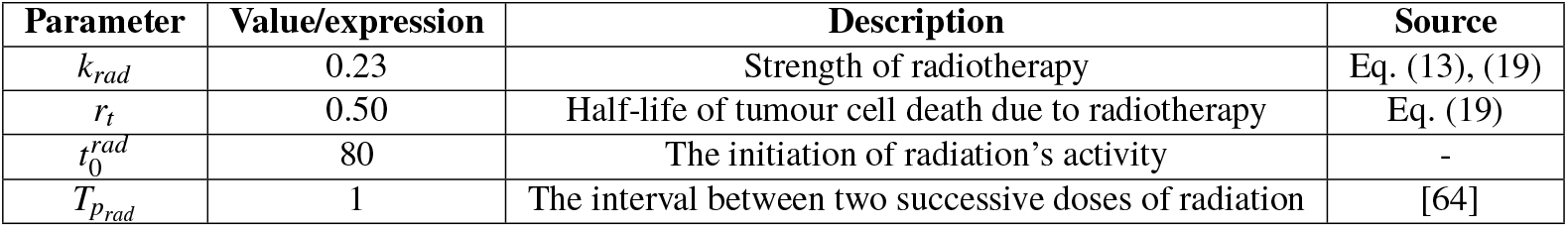
Dimensionless parameter values used in radiotherapy simulations.

### Anti-VEGF drug (Bevacizumab)

The mass balance equation for the anti-vascular endothelial growth factor (anti-VEGF) drug bevacizumab is expressed by Eq. (9). Within this equation, the source term, *s*_*α*_, plays a pivotal role in formulating and understanding the drug’s behavior and effectiveness:

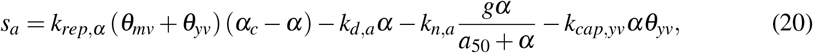

where *k*_*rep,α*_ represents the rate constant for the replenishment of the drug by the vasculature, indicating the efficiency of drug supply to the targeted area; *α*_*c*_ represents the drug concentration in the bloodstream; *k*_*d,α*_ denotes the drug’s decay rate constant in the tissue; *k*_*n,α*_ denotes to the rate constant for drug consumption utilized for neutralizing VEGF. *α*_50_ denotes the drug concentration corresponding to the half maximal rate of VEGF neutralization, commonly referred to as the median effective dose. Finally, *k*_*cap,yv*_ represents the rate constant at which the drug is consumed, inducing endothelial cells apoptosis.

In our proposed model, bevacizumab is administered intravenously during *N* scheduled sessions. If we denote with 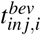 the time of administration *i* (*i* = 1,…, *N*), we model the dynamics of the drug concentration within the bloodstream as follows:

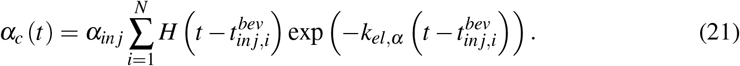

Here, *α*_*in j*_ denotes the drug concentration in the bloodstream at the time of injection, and *k*_*el,α*_ represents the drug elimination rate constant in the bloodstream (determines how quickly the drug is cleared from the body). Eq. (21) can be interpreted as a mathematical model generating a series of exponentially decaying pulses, representing the periodic administration of bevacizumab. In our modeling approach, the time interval between two consecutive sessions is denoted with 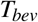, thus 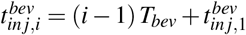.

Including bevacizumab affects the source term of both VEGF (*g*) and young vessels, (*yv*):

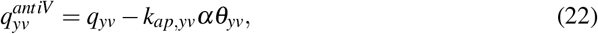

with *k*_*ap,yv*_ being the rate constant of young vessel apoptosis induced by bevacizumab.

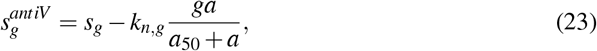

where *k*_*n,g*_ is the VEGF neutralization rate constant.

### Cytotoxic drug (Docetaxel)

To integrate the cytotoxic drug docetaxel into the model, we need first to formulate the mass balance equation for docetaxel adopting the general mass balance equations for chemical species (Eq. (9)). In particular, the source term of docetaxel is formulated as follows:

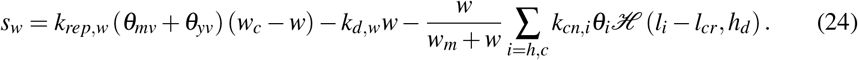

Here, *w* represents the concentration of docetaxel, and *k*_*rep,w*_ denotes docetaxel’s replenishment constant by the vasculature, representing the rate at which the drug is supplied to the tissue. *w*_*c*_ is the concentration of the drug in the bloodstream. The drug decay rate constant in the tissue, denoted as *k*_*d,w*_, indicates the rate at which the drug is metabolized or eliminated from the tissue. The drug consumption rate constants by cancer and healthy cells are denoted as, *k*_*cn,c*_ and *k*_*cn,h*_, respectively. Additionally, *w*_*m*_ signifies the concentration at which the drug consumption rate by cells reaches its half-maximal point. Lastly, *l*_*i*_ represents the proliferation rate of healthy cells (*i* = *h*) and cancer cell (*i* = *c*), and *l*_*cr*_ is a critical value that serves as the criterion for rapidly proliferating cells.

Similarly to bevacizumab, the total number of docetaxel administrations is denoted by *N*. The elimination rate constant of the drug in the bloodstream is denoted by *k*_*el,w*_, and the time of the *i*^*th*^ docetaxel administration is denoted by, 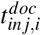. Then, the evolution of docetaxel concentration in the bloodstream, *w*_*c*_ is modelled using:

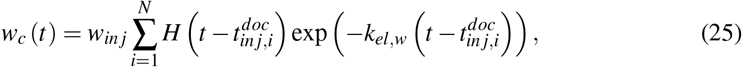

where *w*_*in j*_ is the initial concentration of docetaxel in the bloodstream. If the time interval between two successive administrations is denoted as, *T*_*doc*_, then: 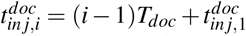.

Furthermore, the inclusion of docetaxel requires appropriate modifications in the source terms for both healthy and cancer cells, denoted as *q*_*h*_ and *q*_*c*_, respectively:

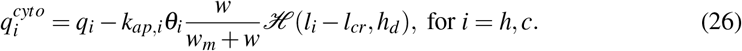

Here, *k*_*ap,i*_ represents the rate constants for the docetaxel induced apoptosis in healthy *k*_*ap,h*_ and cancer cells (*k*_*ap,c*_) alike. We also note that docetaxel specifically targets rapidly proliferating cells [75]. To distinguish between rapidly proliferating cells and those with a lower rate of proliferation, a critical proliferation rate value, *l*_*cr*_, is employed. The smoothened Heaviside function ℋ, activates docetaxel’s killing efficacy (last term of Eq. (26)) for cells with proliferation rate, *l*_*i*_, exceeding the threshold *l*_*cr*_. The proliferation rate is determined by:

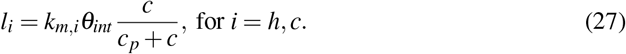

More details are provided in our recent publication [50]. The parameters values associated with chemotherapeutic treatments are provided in the Supplementary Material, chapter II.

## Results

We initiate our investigation by examining how the inclusion of radiotherapy influences the dynamics of a growing tumor as predicted from the proposed mathematical model. Next, we integrate radiotherapy with chemotherapy schedules, specifically in conjunction with both a cytotoxic and anti-VEGF agent. This combination not only broadens the scope of our study but also enables us to investigate potential synergistic effects arising from the simultaneous implementation of these two therapeutic modalities. This approach seeks to capture the intricate interplay treatment modalities and explore the benefits of a multi-faceted therapeutic strategy within the context of our investigation. To provide a comprehensive overview of the various therapeutic protocols employed in our study, we detail the specifics of these treatment regimens in Table 2.

**Table 2.**
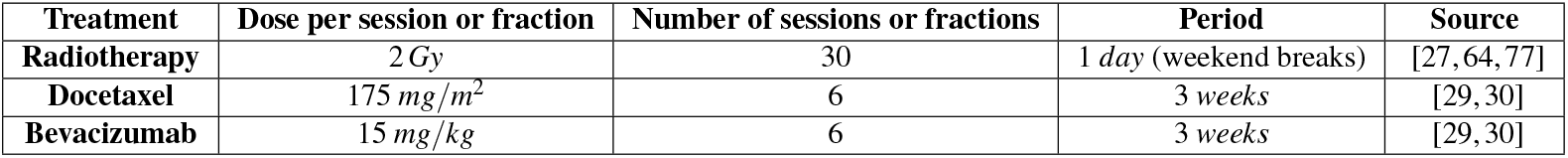
Therapeutic protocols implemented in our current study.

Before presenting the results derived from the application of the aforementioned treatment modalities, it is important to provide a concise overview of the computational mesh and solvers used to address the underlying computational problem. In this study, all computations were performed using Comsol Multiphysics, which is grounded in the principles of the Finite Elements Method (FEM). The infected tissue is modeled by a circular domain with a dimensionless radius, *R*_*tissue*_, set at 30. One unit of length roughly corresponds to *L* = 400 *μm*. This particular length was selected so as to simulate spheroids that are large enough to commence angiogenesis. In addition, one dimensionless unit of time coincides with the time required for a healthy cell’s mitotic division (24 *h*) [76]. The computational domain is discretized using an unstructured mesh generated through the Delaunay Triangulation method, resulting in approximately 70,000 elements and 2,000,000 degrees of freedom. When performing a full simulation using an AMD Ryzen 9 3900X 12-Core Processor for a dimensionless time up to *t* = 650, the required computational time is approximately 70 hours.

### Radiotherapy Simulations

In this section, we investigate the impact of radiotherapy on cancerous tumors. As previously outlined in the introduction, our computational model incorporates an irradiation dose of 2 *Gy* per fraction. The term “fraction” refers to each session during which irradiation is administered to the malignant region. The radiotherapy treatment spans a therapeutic duration of six (6) weeks, ensuring a comprehensive approach to addressing the malignancy [64].

In Fig. 2(a), we depict the evolution of the fraction of the computational domain, *S*, that is covered by cancer cells throughout the simulation following radiotherapy. We refer to this metric as Surface Coverage (*SC*), and is calculated as follows:

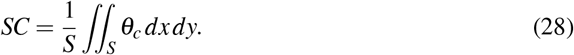

*S* denotes the total surface area of the computational domain, and *θ*_*c*_ is the volume fraction of cancer cells.

**Fig 2.**
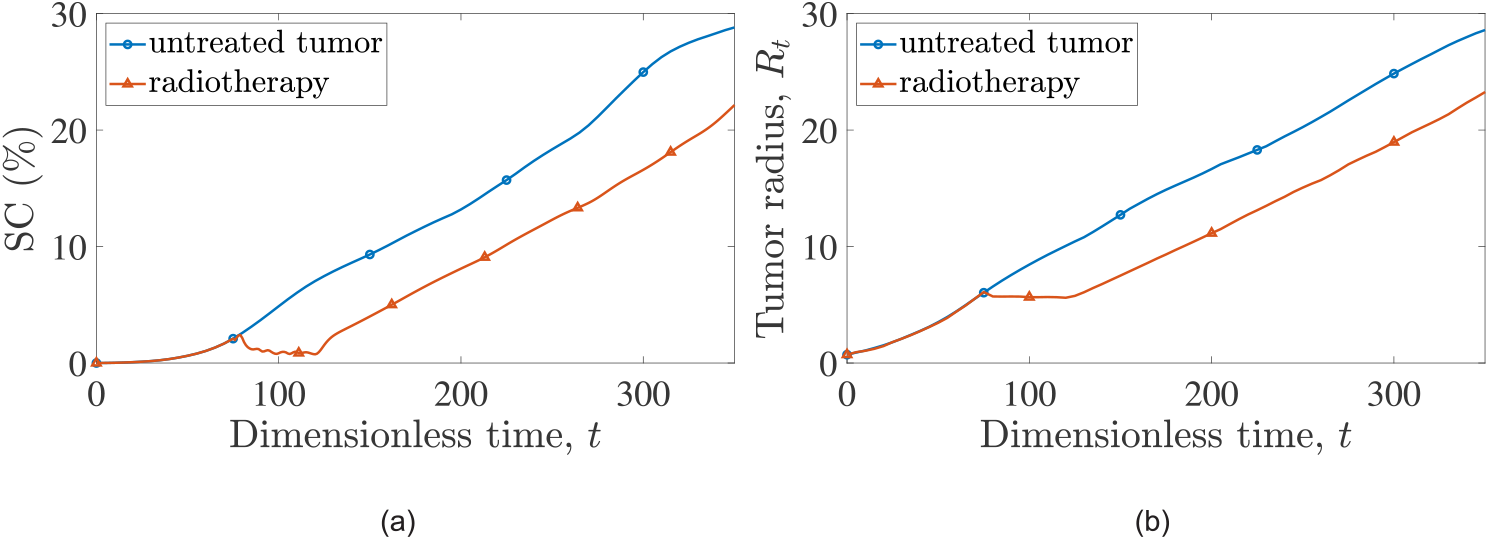
Evolution of (a) cancer cells surface area, *SC* and (b) tumor radius, *R*_*t*_ over time for a simulation of: an untreated tumor (blue line with open circles) and a tumor subjected to radiation treatment starting at 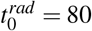 (red line with open triangles).

To assess the efficacy of radiation therapy, we perform a comparison between simulations of untreated tumors and tumors treated with radiotherapy over time. This comparison is performed for both the surface coverage, *SC* (indicative of the total number of surviving cancer cells) and the tumor’s radius as depicted in Fig. 2(b).

The cancer cell surface coverage in the untreated tumor remains higher at all times compared to the irradiated tumor scenario, underscoring the effectiveness of radiotherapy in reducing the number of cancer cells. Notably, one can observe that at the end of all radiotherapy sessions, the *SC* value drops to 0.75% (*t* = 120 dimensionless units), which compares to *SC* = 7% for the treatment-free simulation. Unfortunately, the tumor experiences a relapse shortly after the completion of the treatment regimen, and indicatively, we report that at dimensionless time, *t* = 350, the surface coverage of cancer cells for the treatment-free simulation is 28.8%, whereas for the radiotherapy scenario reduces to 22.1%.

The oscillations during radiotherapy sessions that are depicted in Fig. 2(a) are attributed to the rapid cancer cell decrease during irradiation, which is followed by short relapses (surviving cancer cells start to multiply again when irradiation ceases).

Upon careful examination of Fig. 2(a), one can observe a brief interval, immediately following the completion of all radiotherapy sessions, during which tumor proliferation appears to accelerate before returning to rates akin to those computed for the untreated tumor’s case. This stimulation of cancer growth is attributed to radiation-induced apoptosis within the model. In particular, the material produced by radiation-induced apoptosis enriches the interstitial fluid, thereby providing nourishment to cancer cells. Additionally, the elimination of cancer cells through radiation creates more space for cell expansion and increases oxygen levels within that space.

This observation aligns with experimental observations. Indeed, studies have shown that radiotherapy and cytotoxic chemotherapy-induced apoptosis can stimulate tumor growth [78, 79], partially due to the secretion of pro-apoptotic proteins [80, 81]. Furthermore, stochastic models by Zupanc et al. [82] suggest that the surplus space resulting from radiation treatment can trigger a proliferation response in cancer cells, as they seek to avoid inhibition of cell differentiation caused by encapsulation from neighboring cells.

In Fig. 2(b) we show the tumor radius evolution, *R*_*t*_, which is computed as the average maximum distance at which the cancer cell volume fraction reaches the critical value, *θ*_*c*_ = 0.01. Examining Fig. 2(b), it is clear that radiotherapy effectively stabilizes the tumor radius throughout the treatment sessions. However, once the therapy is completed the surviving cancer cells resume their growth at a rate similar to that observed prior the initiation of therapy.

### Combination of Radiotherapy with Cytotoxic and Anti-VEGF Chemotherapy

In this paragraph, we focus on the application of a synergistic approach involving the concurrent administration of anti-VEGF and cytotoxic chemotherapy drugs, combined with the influential effects of radiotherapy on malignant tumors. The mathematical model detailing the integration of anti-VEGF and cytotoxic agents has been previously presented in [51], where we demonstrated the enhanced efficacy of combined treatment leading to significant tumor shrinkage. Once again in this study, combination therapy was modeled based on typical clinical practices. The therapy scheduling and the doses administered were calculated and non-dimensionalised according to relevant protocols [29, 30]. As mentioned in the previous paragraph, Table2 summarises the schedule and dosage of each treatment. A detailed description of the dosing and its non-dimensionalisation can be found in our previous work [51]. Here, we take a step forward to investigate the effect of radiochemotherapy treatments. For a more comprehensive understanding of the combination therapy model and a detailed table containing all non-dimensionalised parameter values utilized, we refer the reader to Supplementary Material, chapters II and III.

Fig. 3 presents crucial time stamps relevant to the implementation of combination therapy in our model. As illustrated in the figure, we established a shared starting point for the initiation of each therapy at dimensionless time, *t* = 80. At this time, all three treatments are simultaneously administered and subsequently, each unfolds independently. We define the therapy duration to extend until the day of the final therapeutic intervention, specifically the last injection of bevacizumab or docetaxel.

**Fig 3.**
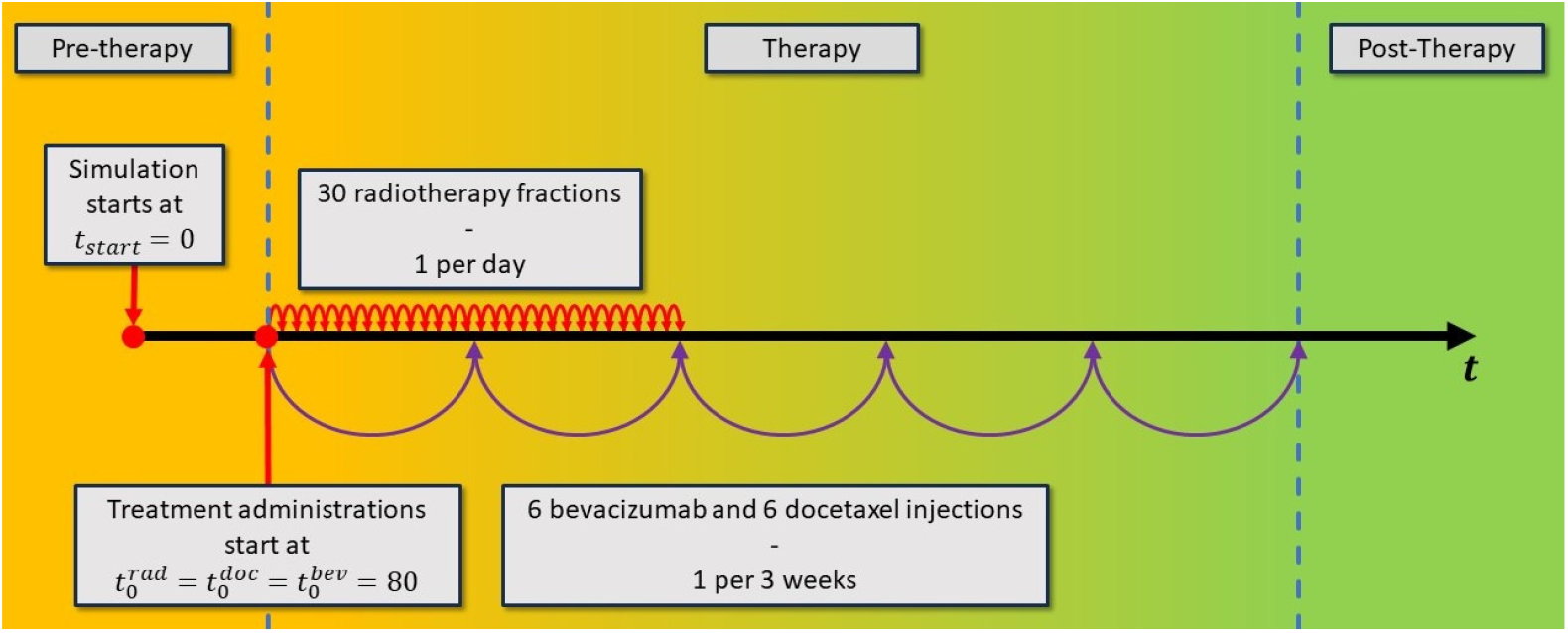
The simulation trajectory of a tumor undergoing treatment with external radiotherapy in combination with bevacizumab and docetaxel.

Similar to the discussion in the previous section, we present *SC* and *R*_*t*_ evolution with time in Fig. 4. To provide additional insight, beyond standalone radiotherapy, and the cumulative impact of all examined therapies, we explore simulations involving tumors treated with docetaxel, bevacizumab, as well as combined treatments using both agents [51]. As shown in both Fig. 4(a) and (b), the combined effect of the three therapeutic agents (combined radiochemotherapy) yields the most favorable outcomes among the considered scenarios. More specifically, referring to Fig. 4(a), it is evident that radiotherapy plays a pivotal role in the therapeutic regiment by hastening treatment results and significantly reducing the cancer cell count. While its long-term impact may not be as pronounced, it imparts a unique benefit by leaving a more suppressed tumor for other agents to manage. Consequently, this results a small tumor throughout the entire simulation course (green curve with open triangles) compared to the one associated with the combined chemotherapy (purple curve with stars).

**Fig 4.**
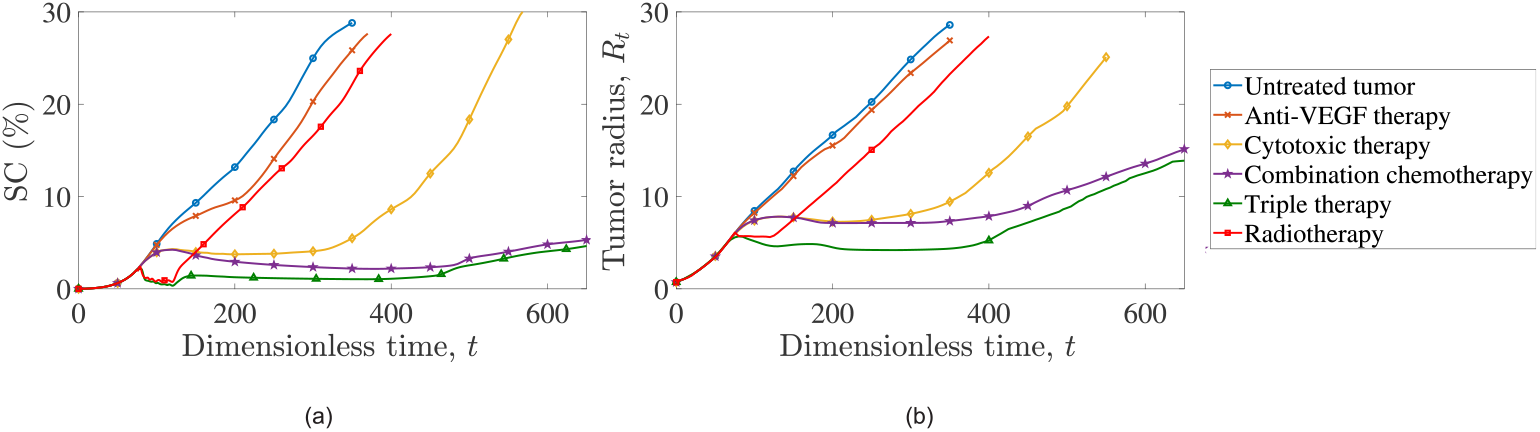
Evolution of (a) cancer cells surface area and (b) tumor radius, ***R***_*t*_ over time for: untreated tumor (blue line with open circles), tumor treated with bevacizumab (orange line with crosses), tumor treated with docetaxel (yellow line with open diamonds), tumor treated with combined chemotherapies (bevacizumab + docetaxel, purple line with open stars), tumor treated with chemoradiotherapy (green line with open triangles) and irradiated tumor (red line with open squares). For all simulations, therapies start at dimensionless time, *t* = 80.

These observations are reinforced by examining Fig. 4(b). As we previously established, radiotherapy has the ability to maintain the tumor’s radius stable for its duration, enabling the co-administration of the other two therapeutic agents to achieve substantial tumor shrinkage. This established baseline has enduring effects, as the tumor requires a considerable amount of time to return to its original size, thanks to the lasting benefits provided by the combination of bevacizumab and docetaxel.

In Fig. 5, we depict snapshots of cancer cell surface distributions for: (i) a therapy-free simulation, (ii) an irradiated tumor, (iii) a docetaxel/bevacizumab combined chemotherapy regimen and (iii) a combined radiochemotherapy simulation. In the first column, the surface distributions depict a growing tumor without any therapeutic interventions. Notably, the three distinct cancer zones (necrotic, quiescent and proliferative) are easily discernible. Moving to the second column, the surface distributions depict a tumor subjected to radiation. Although the three tumor zones remain distinguishable, it is apparent that both the tumor’s radius and the total number of cancer cells have been reduced due to radiotherapy. This tumor exhibits morphological similarity to the untreated tumor, albeit with a delay attributable to the application of radiotherapy.

**Fig 5.**
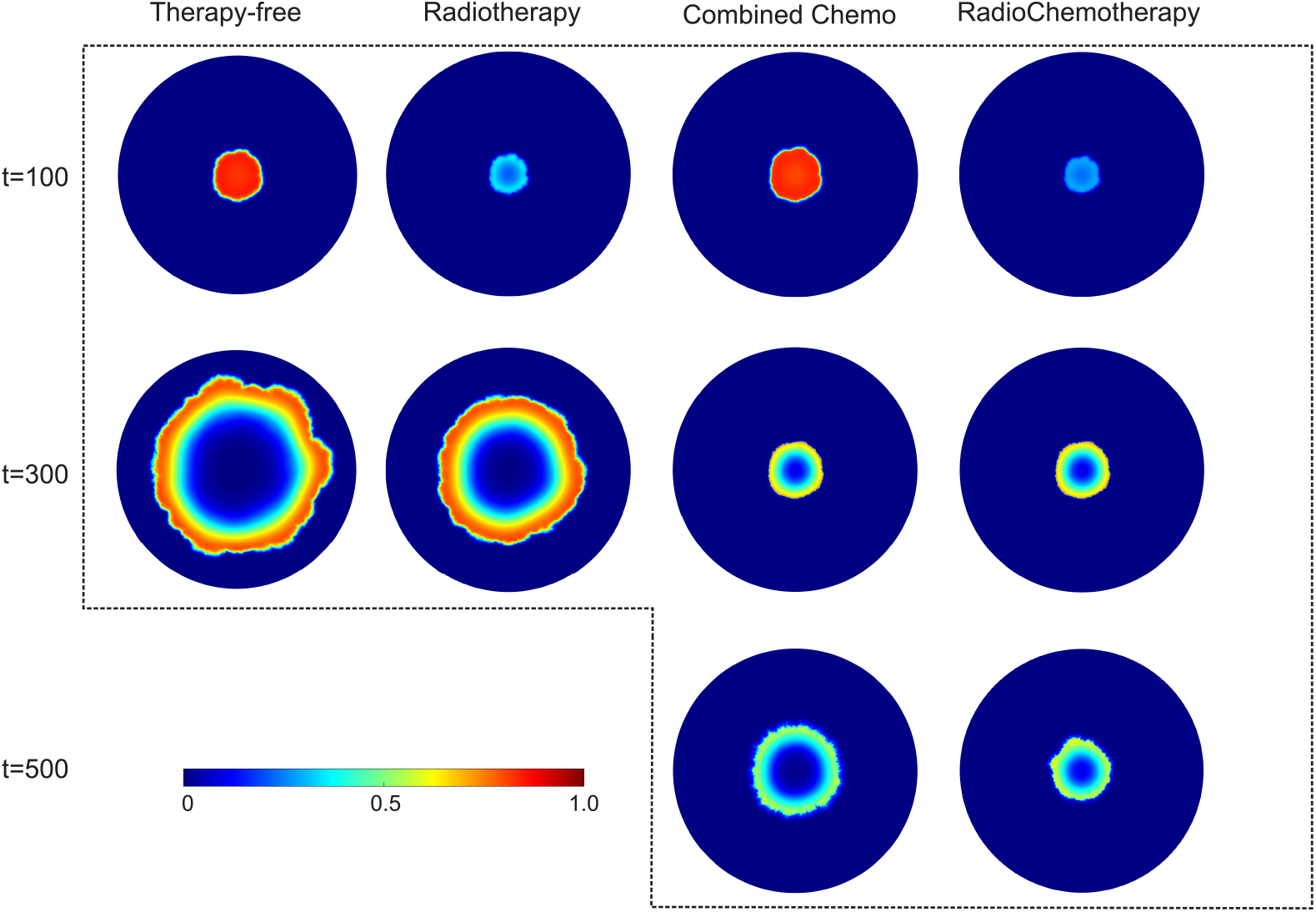
Cancer cell volume fraction distributions, *θ*_*c*_, are depicted in four columns representing different treatment scenarios: (first column) therapy-free, (second column) radiotherapy, (third column) combined chemotherapy (docetaxel and bevacizumab) and radiochemotherapy (fourth column). The first horizontal line depicts cancer cell volume fraction distributions at dimensionless time, *t* = 100, and the second horizontal line for dimensionless time, *t* = 300. By dimensionless time *t* = 500, the therapy-free and radiotherapy simulations show tumors that reach and exceed the computational domain boundaries. In contrast, the distributions for combined chemotherapy and radiochemotherapy at dimensionless time, *t* = 500, reveal tumors with the lowest growth dynamics.

The third column showcases a tumor treated with the bevacizumab/docetaxel combination therapy. Notably, the tumor is suppressed for a significantly prolonged period compared to the tumors in the first two columns. While the short-term effects of radiotherapy are undeniably more potent, the chemical agents appear to hold an advantage in the long term. Finally, the last column presents a tumor treated with all three therapeutic agents (radiochemotherapy). As previously emphasized, triple therapy seamlessly combines the benefits of both radiation therapy -with its immediate results- and the chemical agents -with their sustained efficacy over time. Undoubtedly, this combination promptly yields the highest reduction in cell count and the most effective containment of the tumor radius.

Let us now shift our focus on evaluating the impact of each therapeutic regimen on the healthy cells of the tissue. We introduce a new metric, namely the recession of healthy tissue denoted as *φ* (*t*). This metric quantifies the difference between the initial surface area covered by healthy cells and the corresponding surface are at each time point, *t*:

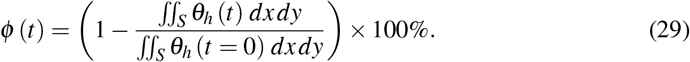

Fig. 6 illustrates *φ* (*t*) for each treatment, including the untreated, therapy-free tumor. The dynamics of healthy tissue recession are influenced by two primary factors. Firstly, the inflammatory properties of cancer play a pivotal role, leading to nutrient deprivation and the displacement of healthy cells. Secondly, the impact of bevacizumab, which binds to VEGF, and impedes the proliferation of endothelial cells, resulting in vascular recession and subsequent tissue damage [23].

**Fig 6.**
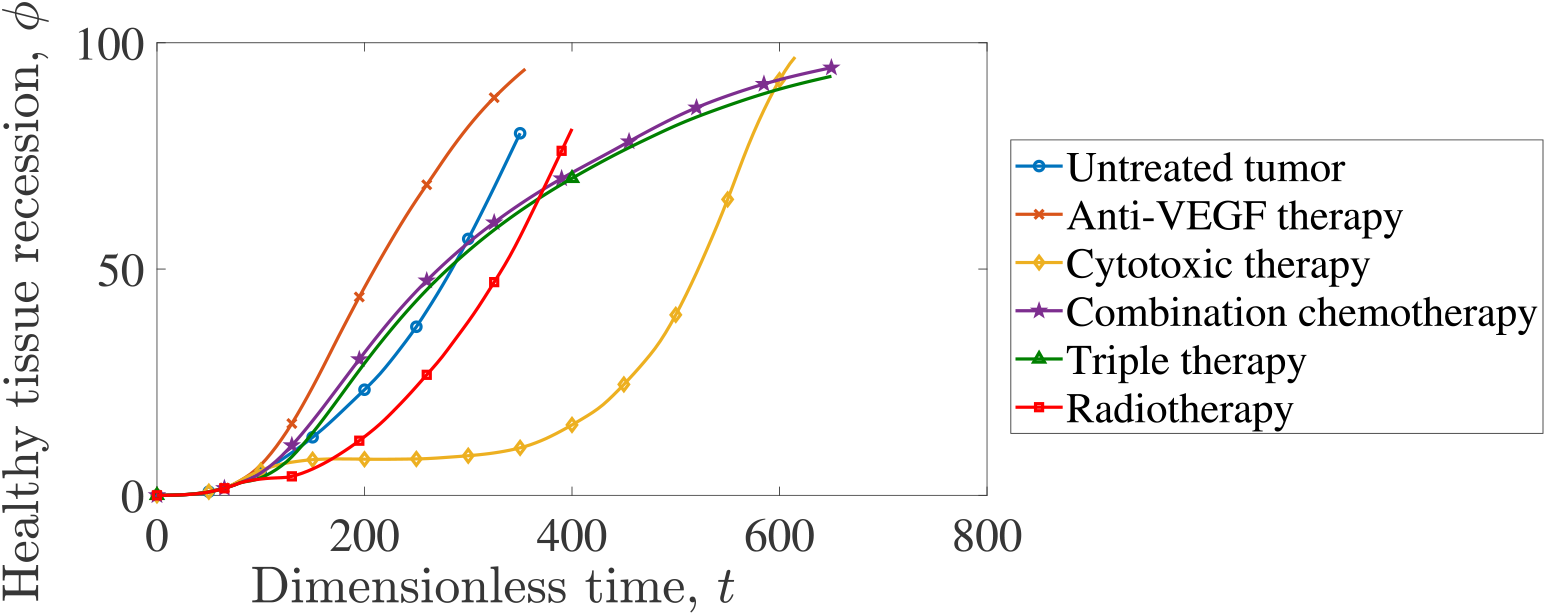
Healthy tissue recession evolution for different therapeutic regimens. We also illustrate the therapy-free simulation with blue line and open circles. Anti-VEGF therapy (crossed orange line) induces the highest healthy cell recession dynamics, suppressing the nutrient supply. The cytotoxic therapy (yellow line with open diamonds) limits only the short-term damage, however not preventing the long term healthy tissue recession. Combination chemotherapy (purple line with open stars), and radiochemotherapy (green line with open triangles) mitigate the negative impact of anti-VEGF therapy.

The therapeutic regimes presented above, effectively manage to mitigate inflammation, thereby safeguarding the surrounding tissue. Radiation therapy aligns with the results depicted in Fig. 2, affording short-term protection to the tissue. In contrast, cytotoxic chemotherapy (docetaxel) significantly limits short-term tissue damage while maintaining a less damaged state for an extended time period. However, at later stages of the simulation the tumor relapse is accompanied by significant healthy tissue damage. Notably, one can observe that the anti-VEGF therapy (bevacizumab) intensifies the healthy tissue damage even when compared to the therapy-free simulation, since its primary functionality is to impede the nutrient supply. These findings are in line with clinical findings. The meta-analysis by Zhao et al. [83] underlines the significant risks of hypertension and proteinuria associated with the use of bevacizumab.

Specifically, hypertension risks were attributed to a few factors. Bevacizumab, functioning as a VEGF inhibitor, induces cell apoptosis and decreases endothelial renewal; consequently, this leads to reduction of capillary density and elevation of the peripheral vascular resistance, thereby impeding blood circulation. In the present model, the reduced vascular density and endothelial renewal are illustrated. Furthermore, Fig. 2 describes exactly the same cell apoptosis induced by bevacizumab.

The combination of bevacizumab with the more favorable docetaxel regimen effectively mitigates the former’s side effects. Even though, the healthy tissue recession grows continuously during the entire course of the simulation, at the final stages of the simulation (*t* ≈ 600) the combination of chemotherapies yields results comparable to cytotoxic monotherapy (*φ* ≈ 90%). Finally, one can observe that the application of chemoradiotherapy reduces the healthy tissue recession compared to the combined chemotherapy regimen.

Returning to the quantification of combination therapy’s efficacy, Fig. 7(a) presents the surface coverage, *SC*, for tumors being treated with a combination of cytotoxic drug, anti-VEGF drug and external radiotherapy. However, in this case, radiotherapy is administered either for half the fractions or, with radiotherapy starting at *t* = 186, after all chemotherapeutic sessions have concluded. Reducing radiation by half significantly mitigates the anti-tumor effect, especially in the later stages of tumor development. Although the tumor’s response remains roughly homologous with the default regime until *t* ≈ 400, it experiences a notable relapse thereafter.

**Fig 7.**
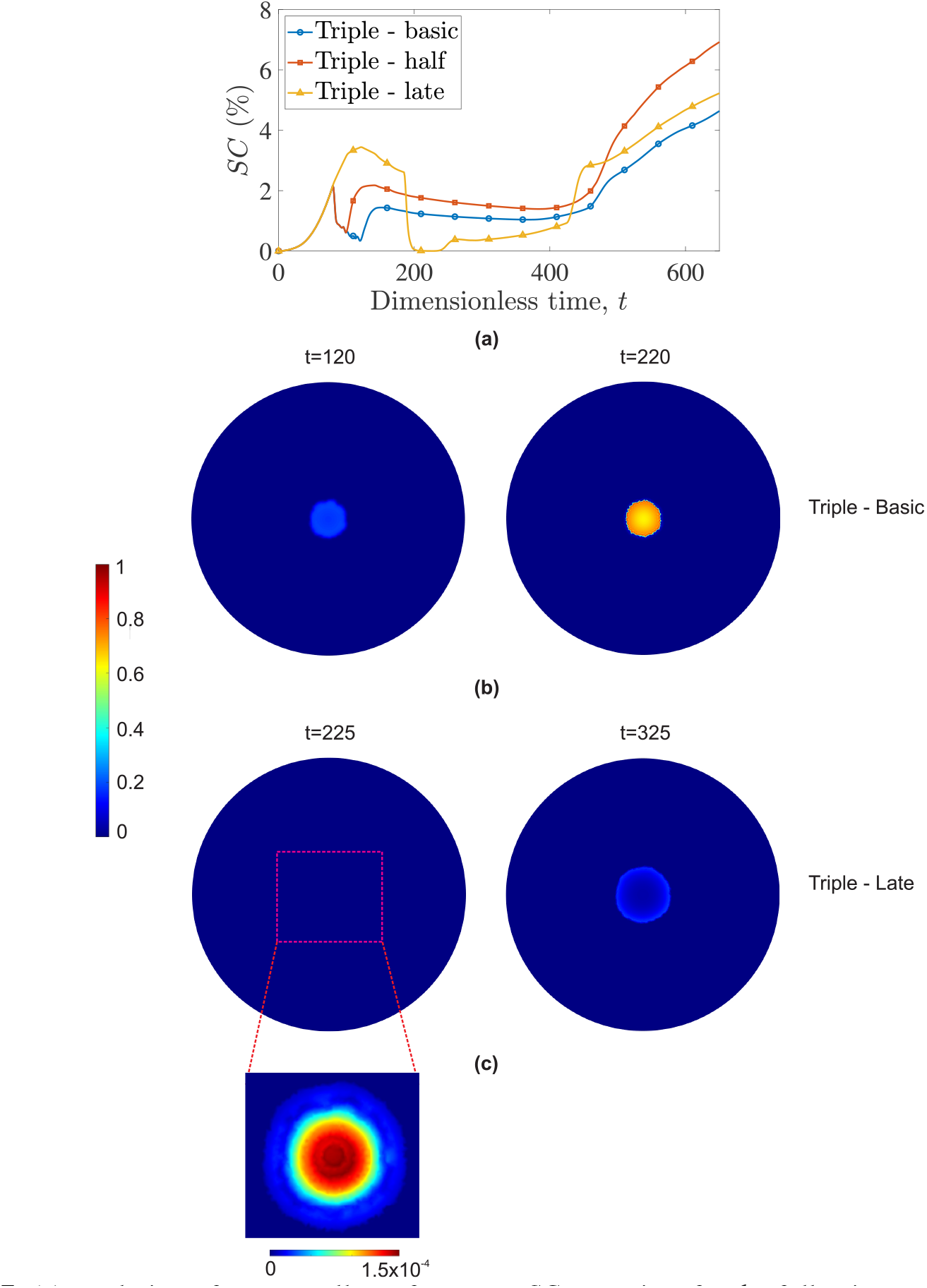
(a) Evolution of cancer cells surface area, *SC*, over time for the following treatments: tumor treated with docetaxel, bevacizumab and radiotherapy all commencing at *t* = 80 (blue curve with circles-Triple basic); the same therapeutic schedule but with half the radiotherapy fractions (red curve with squares - Triple half); and the initial regimen but with radiotherapy beginning after the chemotherapy sessions conclude (*t* = 186) (yellow curve with triangles-Triple late). (b) Surface distribution of cancer cells at *t* = 120 and *t* = 220 for tumor treated with concurrent docetaxel, bevacizumab and radiotherapy. (c) Surface distribution of cancer cells at *t* = 225 and *t* = 325 for tumor treated with delayed radiotherapy. To better display the cancer cell distribution at *t* = 225, the bottom panel corresponds to the distribution of cancer cells using different coloring scale (with a maximum value of 1.5 *×* 10^−4^).

On the other hand, delaying radiation treatment, results in almost total tumor eradication. It is noteworthy that between *t* = 200 and *t* = 230, tumor concentrations universally drop below *θ* = 0.01, rendering *R*_*t*_ (tumor radius) undetectable. However, despite the tumor remaining contained for an extended period, it experiences an aggressive relapse. Admittedly, administering all therapies in parallel leads to a more synergistic action, with docetaxel and bevacizumab targeting a more vulnerable tumor and better managing long-term containment.

Figures 7(b) and (c) depict the surface distributions of cancer cells for concurrent radiotherapy (commencing at *t* = 80) and delayed radiotherapy (commencing at *t* = 186), respectively. The left column shows cancer cell distributions right after the completion of each radiotherapy administration (*t* = 120 and *t* = 225, respectively). The right column presents cancer cell distributions after an interval of 100 dimensionless time points.

Of particular note in this figure is the significant disparity in therapeutic outcomes between the same treatment regimen administered with and without delay. Notably, early/concurrent administration results in the tumor radius remaining relatively stable after a Δ*t* = 100, with a slight increase in cellular density. Conversely, delayed radiotherapy exhausts the tumor, leaving only a small seed capable of spawning a new tumor. By inspecting the tumor at *t* = 225 (left panel of Fig. 7(c)), one can observe that the combined effect of chemo-radiotherapy is particularly effective on the tumor’s periphery. This region, abundant in cellular debris and oxygen, becomes highly conducive to restoration, as evident after Δ*t* = 100. The tumor’s proliferative zone shows signs of restoration, indicating an impending relapse.

Lastly, we are enhancing our understanding of each therapy’s contribution by pairing them up two at a time and comparing their effectiveness. This comparison is illustrated in Fig. 8, where the combinations of docetaxel-radiation, bevacizumab-radiation and bevavicumab-docetaxel are administered in the simulated tumors. It becomes evident that inclusion of docetaxel achieves a long term, tumor suppressive environment. Both combinations that exclude docetaxel exhibit poor long-term efficacy. Furthermore, bevacizumab-radiation leads to immediate tumor relapse after completing the radiation fractions. Interestingly, both radiation and bevacizumab function effectively as adjuncts to cytotoxic chemotherapy. Radiation weakens the tumor while docetaxel concentration builds up, making its target more vulnerable. Bevacizumab, on the other hand, prolongs docetaxel’s residence time in the tissue [51].

**Fig 8.**
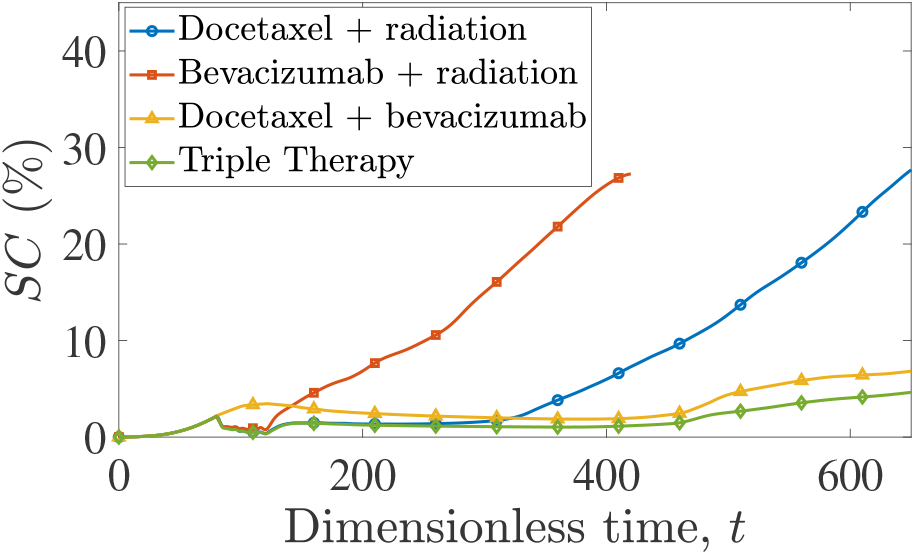
Evolution of cancer cells surface area, *SC*, over time for a tumor treated with: docetaxel - radiation combined therapy (blue line with circles), bevacizumab - radiation combined therapy (red line with squares), docetaxel - bevacizumab combined therapy (yellow line with triangles), and triple therapy (green line with diamonds).

### Radiosensitization

#### Docetaxel-induced Radiosensitization

Certain chemotherapeutic and other agents possess the capability to augment the effects of radiation in tumors, thereby amplifying the DNA damage caused by radiation [84]. Taxanes, such as docetaxel or paclitaxel, are among these agents. Their mechanism of action entails arresting the cell cycle during the G2/M phase, a phase when cells are highly radiosensitive [85–88]. Recent clinical trials have corroborated this finding by demonstrating promising results regarding the inclusion of docetaxel as a radiosensitizer in patients ineligible for cisplatin-based chemoradiation. Specifically, this therapeutic regimen has shown improvements in both overall survival and progression-free survival among the studied patients [89].

We formulate radiosensitization in our model, by modifying the kinetic term describing the cellular death caused by external radiotherapy, as presented in Eq. (10):

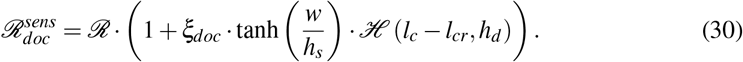

*ξ*_*doc*_ serves as the radiation’s potency amplifier due to the presence of docetaxel; *h*_*s*_ is an appropriately selected smoothing parameter determined by doectaxel’s concentration. According to Eq. (30), the radiation’s potency is amplified with increasing values of docetaxel’s concentrations, *w*. In addition, the radiotherapy-induced killing rate is amplified only for highly proliferative cancer cells the cycle of which is disrupted by the cytotoxic drug. Targeting of highly proliferative cancer cells is modeled utilizing the Heaviside function, ℋ .

Figure 9 highlights the enhanced efficacy of triple-combined therapy, considering the phenomenon of radiosensitization. In particular, we examine the impact of radiosensitization in two cases: (a) when all treatments are administered simultaneously, and (b) when radiation follows the completion of chemotherapy. In both scenarios, radiosensitization enhances the efficacy of the administered therapies, as evidenced by consistenly lower *SC* values. Particularly in the case of delayed radiation regimen, radiosensitization extends the duration during which the tumor approaches eradication and mitigates the aggressiveness of tumor relapse.

**Fig 9.**
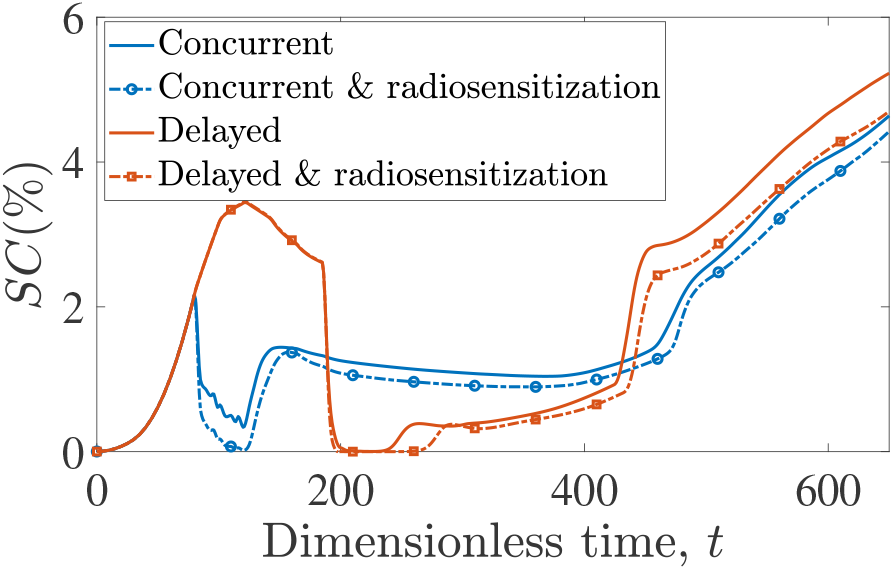
The evolution of cancer cell surface area, *SC*, over time for tumors treated with docetaxel, bevacizumab and radiation all starting at *t* = 80 (concurrent) with and without considering docetaxel-induced radiosensitization is shown with blue and blue dotted curves, respectively. The same regimen but with radiotherapy starting after the completion of chemotherapy sessions (delayed) is depicted with red curves when neglecting radiosensitization, and with red dotted curves incorporating radiosensitization.

### Oxygen Radiosensitization

Another important parameter influencing the effectiveness of radiation is the level of oxygenation in the targeted cells. Oxygen acts as a potent radiosensitizer, significantly boosting the ability of radiation to eradicate cancer cells [55]. In fact, hypoxic cells can exhibit resistance to radiation therapy, contrary to well-oxygenated cells [90, 91].

To incorporate the impact of oxygen-induced radiosensitization into our model, we modify the existing kinetic term, ℛ. In contrast to Eq. (30), where the initial term remained unaltered and was multiplied by an additional factor, here, the initial *k*_*rad*_ is substituted to generate a term with an equivalent effect during *standalone/monotherapy* radiation treatment:

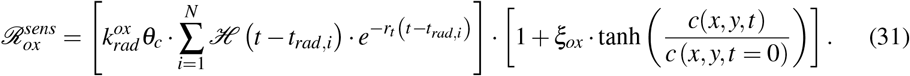

The updated kinetic term for cellular death induced by external radiation integrates its heightened efficacy against well-oxygenated cells. The term tanh 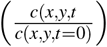 ensures that the death rate constant, 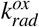, is amplified with increasing oxygen concentration relative to its initial value.

To ensure an equivalent outcome between the kinetic terms ℛ and 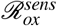 when *standalone* radiotherapy is administered, we derived a new rate constant 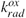, so as to ensure that *k*_*rad*_ and 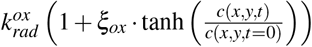 produce a comparable killing effect. For a more comprehensive description of the derivation steps for the value of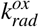, we refer the reader to Chapter IV of SI.

To underline the impact of oxygen on radiation therapy, we present the (average) radial distribution of cancer cells at *t* = 120 (immediately after the last fraction has been administered) in Fig. 10 for tumors treated solely with radiation therapy, both with and without consideration of oxygen sensitization. As previously mentioned, the overall radiation effect is comparable for both scenarios, since both the cancer cell surface coverage, *SC*, and the tumor’s radius, *R*_*t*_ were maintained at approximately equal levels. However, the spatial distribution of cancer cells exhibits notable differences between the two cases, as depicted in Fig. 10. To enhance readability, we opted to present the average cancer cell volume fraction for each azimuthal angle across the tumor’s radius (considering an approximately circular shape of the growing tumor). Moreover, we rescaled the volume fractions for each tumor 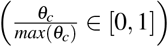, to provide a clearer representation of how cancer cells are relatively allocated within the tumor in either case of irradiation. Indeed, when considering calculations that neglect radiosensitization phenomena as the default, Fig. 10 shows the significant impact of such phenomena on the tumor’s morphology. In other words, rather than uniformly affecting cancer cells, we observe that -in comparison to our established default scenario-cells in well-oxygenated regions near to the tumor’s periphery exhibit a more pronounced reduction, while the interior of the tumor shows up to a 50% lesser degress of effect.

**Fig 10.**
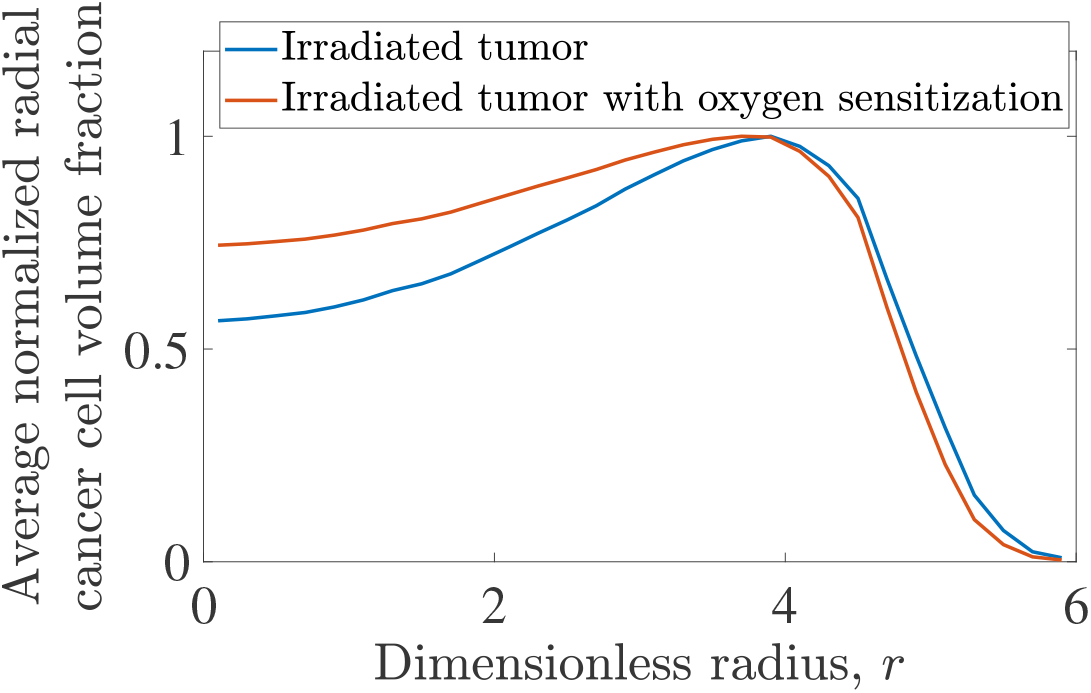
Average cancer cell volume fraction radial distribution, normalized with maximum cancer cell volume fraction for tumors irradiated with external radiotherapy: blue curve corresponds to simulations neglecting oxygen sensitization. The red line illustrates the corresponding distribution incorporating oxygen sensitization. Both distributions are computed at dimensionless time, *t* = 120, following the last fraction of administered radiotherapy.

We now delve into examining the incorporation of oxygen radiosensitization when administering triple-combined therapy to treat tumors. In Fig, 11(a)&(b), simulations are presented for tumors undergoing a combination of chemotherapy and radiotherapy, with consideration given to either chemo- or oxygen-induced radiosensitization effects on tumor cells. Of particular interest in these findings, unlike radiosensitization from docetaxel, which confers benefits regardless of the therapeutic schedule employed, the reliance of radiation treatment on cellular oxygenation underscores the importance of timing in its administration for optimal outcomes. Fig. 11(a) shows the comparison between simulations with and without consideration of radiosensitization effects when treatment commences at earlier stages of tumor growth (*t* = 80), whereas Fig. 11(b) shows the comparison for the scenario in which therapy is administered at later stages of tumor growth. As depicted in Fig. 11(b), it is evident that as the tumor progresses and depletes tissue oxygen, radiation efficacy diminishes as cells become radio-resistant in hypoxic conditions. This is verified by looking at Fig. 11(c) and (d) which present the surface distributions of oxygen in the domain when radiotherapy starts. For Fig. 11(c), the selected time is *t* = 80, corresponding to concurrent radiochemotherapy’s commencement and for Fig. 11(c) *t* = 186, presenting the point in time delayed radiotherapy begins. It it clear that delayed radiotherapy encounters a more oxygen-depleted system, ergo less radiosensitive.

**Fig 11.**
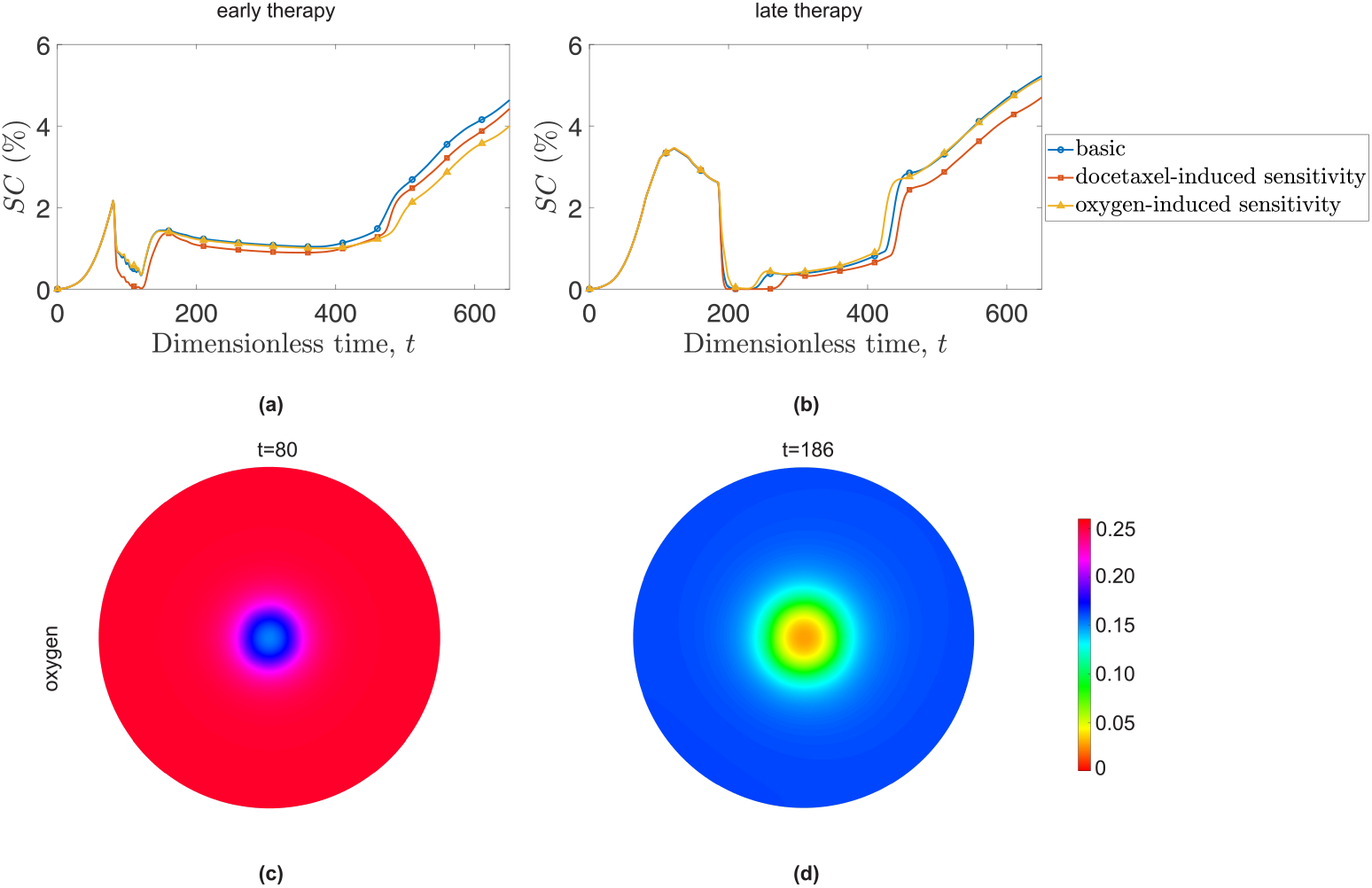
Impact of docetaxel and oxygen-induced radiosensitization on early (left column) and late therapeutic schedules (right column). The first horizontal line ((a) & (b)) illustrates the evolution of cancer cells surface area, *SC*, over time for tumors treated with triple-therapeutic schemes. The blue lines with circles show the evolution of *SC* when radiosensitization is not incorporated in the model, the squared red lines represent *SC* for the case of docetaxel-induced radiosensitization and the yellow lines with triangles illustrate *SC* for the scenario of oxygen-induced radiosensitization. The second horizontal line ((c) & (d)) displays the distribution of oxygen concentration when radiation starts at *t* = 80 (early therapy) and at *t* = 186 (late therapy).

The new parameters associated with radiosensitization are presented in Table 3.

**Table 3.**
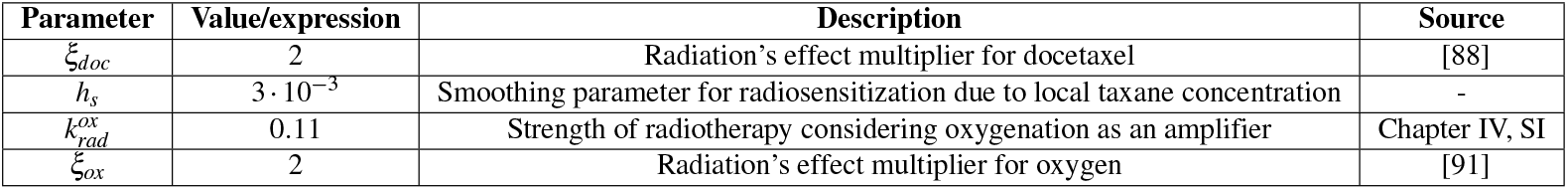
Dimensionless parameter values used in radiotherapy simulations with radiosensitization.

## Conclusion

Our study centers on modeling contaminated tissue as a highly viscous and multiphase fluid. Utilizing a continuous model framework, we employ mass and momentum balance equations solving them with the Finite Elements Method (FEM) through the commercial software Comsol Multiphysics ^®^to ensure both accuracy and efficiency.

A distinctive feature of our model is its versatility, seamlessly integrating various therapeutic agents, including cytotoxic chemotherapy (docetaxel), and anti-VEGF (bevacizumab), with external radiation. As stated in the Introduction, this is the first study integrating all three therapies (cytotoxic, anti-angiogenic and radiation) in one model and monitoring their interplay and short vs long-term intricacies of this schema.

The model showcases the synergistic effects of combined radiochemotherapy, elucidating the intricate interplay between the immediate action of radiation and the long-term effectiveness of chemotherapy. That synergistic effect between therapeutic modalities is further explored after taking into consideration the phenomena of oxygen and taxane induced radiosensitization.

Moreover, the presented model aligns with experimental observations, supporting the effectiveness of the radiation-taxane combination and the inclusion of bevacizumab, as reported in studies such as [92, 93]. In clinical practice, taxane-based chemotherapy is combined with radiation for non-small cell lung cancer instances [27] and gastroesophageal carcinomas [28] for a duration of 5 to 6 weeks. For the treatment of non-small cell lung cancer in particular [27], the radiation treatment duration is six weeks and a total dosage of 60 *Gy* - this is exactly the treatment adopted in our model. On top of that, our model manages to capture the accelerated tumor growth that is experimentally reported after the completion of radiotherapy [78, 79, 82]. Lastly, we managed to showcase the enhanced cancer cell killing efficacy of radiation therapy when combined with docetaxel [85–88] and on well oxygenated targets [90, 91].

In terms of validation protocols, our deliberate decision to deviate from the specific protocol for docetaxel inclusion serves the purpose of maintaining a consistent frame of reference for radiotherapy inclusion. This ensures a fair comparison between treatment schemas (bev-doc and rad-bev-doc).

While our research has made significant strides in exploring combination therapy and understanding short- and long-term tumor responses, there is still a significantly large landscape to explore. Combinations involving platinum-based drugs, commonly used in conjuction with bevacizumab and taxanes, warrant further investigation [29, 30, 94, 95]. Additionally, non-classic cytotoxic chemotherapy forms, such as immunotherapy [96] (e.g., virotherapy, T-cell therapy, macrophage phenotype targeting), hold promise and can be seamlessly integrated into our model.

It is crucial to report that this study presents only one radiochemotherapy scheme, without delving into optimizing parameters, especially considering the intricate scheduling of different therapeutic factors. Our previous work [51] underscored the pivotal role of therapy scheduling in treatment success. Given the inclusion of three therapeutic factors in our computational experiments, leveraging Gaussian optimization and other machine learning tools can systematically optimize scheduling parameters to enhance treatment success.

## Acknowledgments

The authors acknowledge the support from the Basic Research Program, NTUA, PEVE (No. 65232000) for this work.

